# A non-toxic equinatoxin-II reveals the dynamics of sphingomyelin in the cytosolic leaflet of the plasma membrane

**DOI:** 10.1101/2023.11.10.566659

**Authors:** Toshiki Mori, Takahiro Niki, Yasunori Uchida, Kojiro Mukai, Yoshihiko Kuchitsu, Takuma Kishimoto, Asami Makino, Toshihide Kobayashi, Hiroyuki Arai, Yasunari Yokota, Tomohiko Taguchi, Kenichi G.N. Suzuki

## Abstract

Super-resolution microscopic observation of a novel non-toxic sphingomyelin probe revealed the formation of dynamic small domains including sphingomyelin and cholesterol in the cytosolic leaflet of living cell plasma membranes.

**Abstract:** Sphingomyelin (SM) is a major sphingolipid in mammalian cells. SM is enriched in the extracellular leaflet of the plasma membrane (PM). Besides this localization, recent electron microscopic and biochemical studies suggest the presence of SM in the cytosolic leaflet of the PM. In the present study, we generated a non-toxic SM-binding variant (NT-EqtII) based on equinatoxin-II (EqtII) from the sea anemone *Actinia equina*, and examined the dynamics of SM in the cytosolic leaflet of living cell PMs. NT-EqtII with two point mutations (Leu26Ala and Pro81Ala) had essentially the same specificity and affinity to SM as wild-type EqtII. NT-EqtII expressed in the cytosol was recruited to the PM in various cell lines. Super-resolution microscopic observation revealed that NT-EqtII formed tiny domains that were significantly colocalized with cholesterol and N-terminal Lyn. Meanwhile, all the examined lipid probes including NT-EqtII underwent apparent fast simple Brownian diffusion, exhibiting that SM and other lipids in the cytosolic leaflet rapidly moved in and out of domains. Thus, the novel SM-binding probe demonstrated the presence of the raft-like domain in the cytosolic leaflet of living cell PMs.

## Introduction

SM is a major sphingolipid in mammalian cells and enriched specifically in the extracellular leaflet of the PM. Because of its high content of saturated acyl chains, SM, along with cholesterol and glycosphingolipids, is suggested to form lipid nanodomains or “lipid rafts” (Lingwood and Simons, 2010). Lipid rafts promote the accumulation of distinct sets of proteins, including ones anchored in the extracellular leaflet by a glycosylphosphatidylinositol moiety (Paulick and Bertozzi, 2008) and ones anchored in the cytosolic leaflet by lipid modification (Prior et al., 2003).

At present, there are a few SM-binding proteins available to visualize SM in cells. Lysenin is a SM-binding protein toxin that was isolated from the coelomic fluid of the earthworm *Eisenia foetida* (Yamaji et al., 1998). Lysenin binds to SM when SM is clustered (Ishitsuka et al., 2004). EqtII is another SM-binding protein toxin isolated from the sea anemone *Actinia equina* (Bakrac et al., 2008) and preferentially binds to dispersed SM (Makino et al., 2015). EGFP-tagged recombinant lysenin and EqtII have been used to detect SM in the PM and intracellular organelles (Yachi et al., 2012). A study with a freeze-fracture replica electron microscopy with recombinant lysenin suggested the presence of SM in the cytosolic leaflet of the PM (Murate et al., 2015). This notion is further supported by a biochemical study, in which the lipid species in the extracellular or the cytosolic leaflet were quantitated (Lorent et al., 2020). Targeting the cytosolically expressed bacterial sphingomyelinase (bSMase) to the PM increased the levels of cellular ceramide, also suggesting the presence of SM in the cytosolic leaflet of the PM (Sakamoto et al., 2019).

In the present study, we generated a non-toxic variant of EqtII (NT-EqtII) so as to overcome the problem of cytotoxicity of EqtII when expressed in the cytosol. With this new probe in our hands, we could characterize the dynamics of SM in the cytosolic leaflet of the PM in living cells for the first time.

## Results

### Generation of a non-toxic EqtII variant

In order to visualize SM in the cytosolic leaflet of the PM, we sought to generate the EqtII variant that lacks pore-forming activity. The pore formation by EqtII is suggested to proceed through a series of sequential processes: monomeric EqtII binding onto the surface of SM-containing membrane, its *N*-terminal helix insertion into membrane, and oligomerization to generate a final pore (Rojko et al., 2013). Given that the oligomerization of the helix appears to be critical for the cytolytic activity that is coupled with the formation of the proteinaceous pore, we attempted to generate EqtII variants with mutations in the helix or in its vicinity. Leu26 is positioned in the *N*-terminal helix (Subburaj et al., 2015). By molecular dynamics simulation, Pro81 in the loop between β5 and β6 was predicted to be buried in the membrane (Weber et al., 2015). Please be noted that the mutation of Val22Trp, Val8Cys/Lys69Cys, or Tyr108Iso of EqtII yielded non-toxic EqtII (Bakrac et al., 2010; Deng et al., 2016) and that these variants have been used to monitor the exposure of SM in the cytosolic leaflet of organelles.

Fortuitously, we found that the simultaneous mutations of these two amino acid residues to Ala abolished the cytotoxicity of EqtII. Three recombinant proteins tagged with His-tag and EGFP [His-EqtII-EGFP, His-EqtII (L26A/P81A)-EGFP, and His-EqtII (L26A/P81A/Y113A)-EGFP] were expressed in *Escherichia coli* and purified. The mutation of Tyr113 to Ala was reported to diminish the SM-binding ability of EqtII (Bakrac et al., 2010). We examined their cytotoxicity with three conventional assays: lactate dehydrogenase (LDH) release assay, WST-8 assay, and hemolysis assay. As shown (Fig. 1A), His-EqtII (L26A/P81A)-EGFP and His-EqtII (L26A/P81A/Y113A)-EGFP did not exert the cytotoxic effect at their concentrations up to 10,000 ng/ml. In contrast, His-EqtII-EGFP showed the effect on cellular viability and LDH release at 1,000 ng/ml and on hemolysis at 100 ng/ml.

**Fig. 1.**
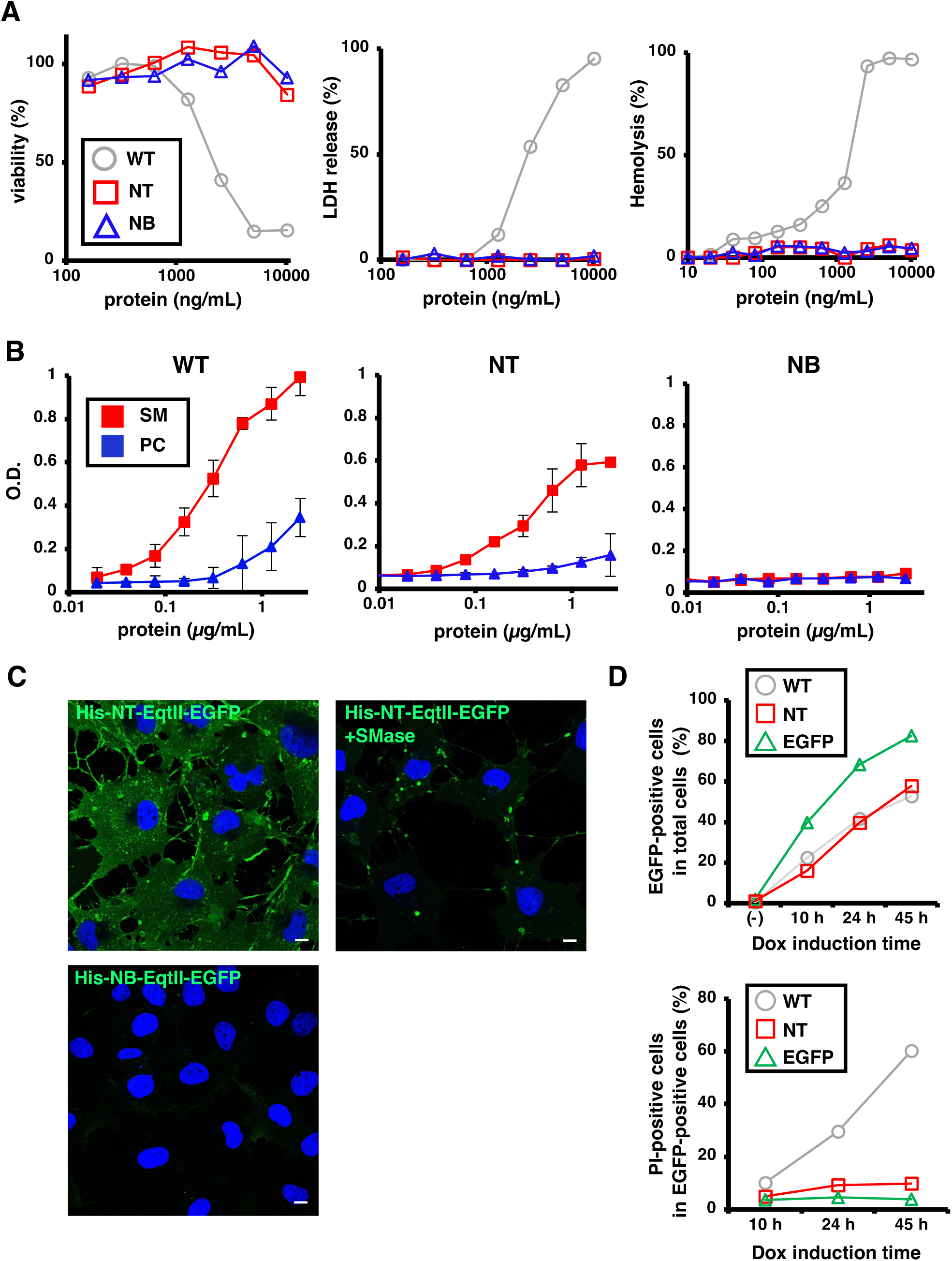
A dual mutation of EqtII abrogates its pore-forming cytotoxicity. (A) (Left and the middle panels) COS-1 cells were treated with recombinant EqtII proteins [WT, His-WT-EqtII-EGFP; NT, His-NT-EqtII (L26A/P81A)-EGFP; or NB, His-NB-EqtII (L26A/P81A/Y113A)-EGFP] at the indicated concentration for 30 min. Cell viability and cytotoxicity were then determined with WST-8 assay and LDH release assay, respectively. (Right panel) Mouse erythrocytes were treated with each protein at the indicated concentrations for 30 min. Hemolysis was then determined by measuring the absorbance at 600 nm. (B) Binding of His-EqtII proteins to SM or PC was determined by ELISA with anti-His-tag antibody. Data represent means ± standard deviations of three (WT) or two (NT and NB) experiments. (C) COS-1 cells were fixed and stained with 1 µg/mL His-NT-EqtII-EGFP or His-NB-EqtII-EGFP. Bacterial SMase was added to the cells 30 min before fixation. Nuclei were also stained with DAPI (blue). Scale bars, 10 µm. (D) COS-1 cells that stably express WT-EqtII-EGFP, NT-EqtII-EGFP, or EGFP in the cytosol in a doxycycline (Dox)-inducible manner were treated with Dox at 1 µg/mL. Cells were then stained with propidium iodide (PI) and analyzed with flow cytometry. Percentages of EGFP-positive cells in total cells (upper panel) and PI-positive cells in EGFP-positive cells (lower panel) are shown at 10 h, 24 h, or 45 h after Dox induction.

Next, we examined whether His-EqtII (L26A/P81A)-EGFP retained the SM-binding ability. His-EqtII (L26A/P81A)-EGFP bound preferentially to SM over PC at its concentration up to 2.5 μg/ml (Fig. 1B). Its affinity to SM was slightly lesser than the affinity of His-EqtII-EGFP to SM. As expected, His-EqtII (L26A/P81A/Y113A)-EGFP did not bind to SM (Bakrac et al., 2010). The specific affinity of EqtII (L26A/P81A) to SM in biomembranes was confirmed by cell staining experiments. When added to the culture medium, His-EqtII (L26A/P81A)-EGFP stained the PM of COS-1 cells, but not that of COS-1 cells pretreated with bSMase (Fig. 1C). His-EqtII (L26A/P81A/Y113A)-EGFP did not stain the PM. Thus, we refer to EqtII (L26A/P81A) and EqtII (L26A/P81A/Y113A) as NT-EqtII (<u>n</u>on-toxic EqtII) and NB-EqtII (<u>n</u>on-lipid <u>b</u>inding-EqtII), respectively.

Lastly, we examined whether NT-EqtII could be expressed in the cytosol. We generated COS-1 cells that stably express WT-EqtII-EGFP, NT-EqtII-EGFP, or EGFP in the cytosol in a doxycycline (Dox)-inducible manner. Propidium iodide (PI) was used as a cell death indicator. After the Dox-induction, the intensities of fluorescence of PI and EGFP in individual cells were quantitated by flow cytometry. PI-and EGFP-double-positive cells were regarded as dead cells. During the Dox-induction up to 45 h, the percentage of cells positive with EGFP signal increased gradually (Fig. 1D). After 45-h induction, the percentage of PI-positive cells in cells expressing WT-EqtII-EGFP was about 60% [31.7/(31.7+21)x100=60.2] (Fig. S1), in line with the cytotoxic effect of WT-EqtII. In contrast, the percentage of PI-positive cells in cells expressing NT-EqtII-EGFP dropped to less than 10% [5.66/(5.66+52.1)x100=9.8]. These results suggested that NT-EqtII could be expressed in the cytosol with less cytotoxicity.

### Single-molecule observation of NT-EqtII revealed the presence of SM in the cytosolic leaflet of the PM

Next, we examined the existence of SM in the cytosolic leaflet of the PM of living immortalized mouse embryonic fibroblasts (iMEFs). iMEFs were transiently transfected with NT-EqtII-HaloTag7, and NT-EqtII-HaloTag7 was visualized by a HaloTag®ligand SF650B. Total internal reflection microscopy (TIRFM) revealed numerous individual fluorescent spots of NT-EqtII-HaloTag7-SF650B on the PM (Fig. 2A-C). We confirmed that single-fluorescent molecules exclusively located in Golgi (TGN38-HaloTag7-SF650B) or ER (mEos4b-STING) were hardly observed by TIRFM. In contrast, the number of puncta was drastically reduced in iMEFs established from sphingomyelin synthase (SMS) 1 and 2-double knockout (DKO) mice. For proper comparison, the number of the puncta was normalized relative to the expressed amount of NT-EqtII. Furthermore, the number of the puncta was also reduced significantly by the expression of bSMase tethered to the cytosolic leaflet of the PM by Ras farnesylation sequence (Sakamoto et al., 2019). The catalytically inactive bSMase mutant (D322A/H323A) did not interfere with the number of the puncta. Treatment of cells for 5 h with 200 μM D609, the inhibitor of SMS, also reduced the number of the puncta. These results elucidated the localization of SM in the cytosolic leaflet of the PM of living iMEFs.

**Fig. 2.**
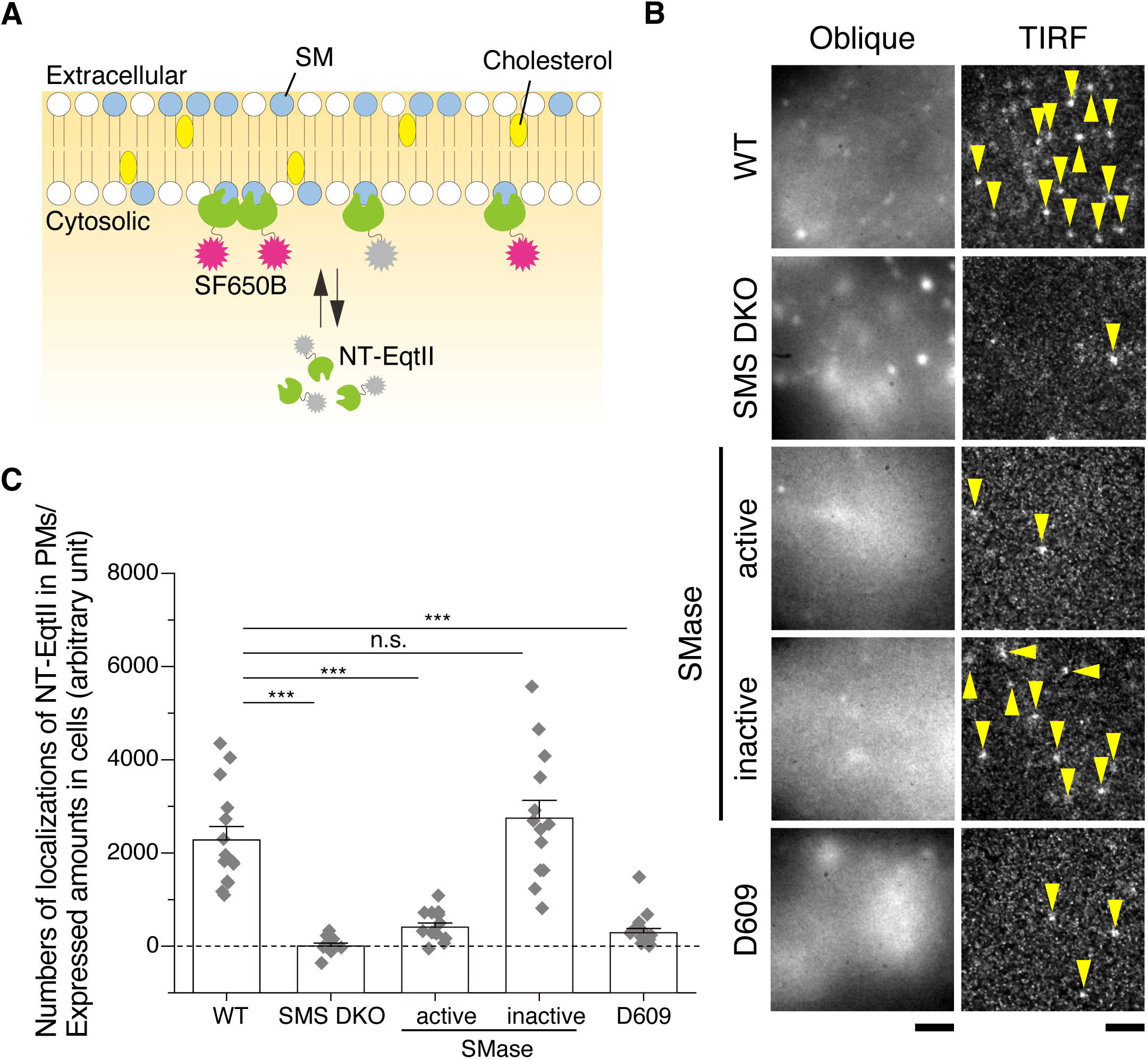
NT-EqtII exhibits a specific affinity for sphingomyelin located in the cytosolic leaflet of the living iMEF cell PM. (A) Schematic representation of NT-Eqt II observation in the cytosolic leaflet of the cell PM. (B) Fluorescent images of NT-EqtII-HaloTag7 labeled with SF650B in various cellular contexts, encompassing wild-type (WT) iMEFs, SMS1 and 2-DKO iMEFs, iMEFs expressing WT bSMase or its catalytically inactive mutant harboring Ras farnesylation sequence by which is anchored to the cytosolic leaflet of the PM, along with iMEFs subjected to D609 treatment. The images were acquired by oblique illumination (left) and total internal reflection fluorescence (TIRF) microscopy (right). Quantification of NT-EqtII-HaloTag7 expressed in cells was performed by evaluating fluorescence intensity via oblique illumination. The content of the SM probe in the cytosolic leaflet of the cell PM was quantified by measuring the numbers of single spots (yellow arrowheads) by TIRFM. Scale bars, 2μm. (C) The quantitated levels of NT-EqtII-HaloTag7 that exhibited binding to the cytosolic leaflet of the cell PM, were subsequently normalized with respect to the expression levels in the cells. The presented error bars symbolize standard errors. *** and n.s. indicate significant (p<0.001) and not significant differences (p>0.05), respectively, using Welch’s T-test.

We sought to determine whether the presence of SM in the cytosolic leaflet of the PM was a general feature in mammalian cells, and thus performed single-molecule observation of NT-EqtII-HaloTag7-SF650B in 14 other cell lines. As shown (Fig. 3), many fluorescent spots of NT-EqtII were observed in 7 cell lines (A549, B16, BxPC3, COS-1, HEK293, PC3, and PZ). The number of the puncta in RBL-2H3 cells was also larger than that of the other 6 cell lines (CHO-K1, HeLa-MZ, HS-5, MRC-5, T24, and WI38). These results showed that a variety of mammalian cell lines expressed SM in the cytosolic leaflet of the PM.

**Fig. 3.**
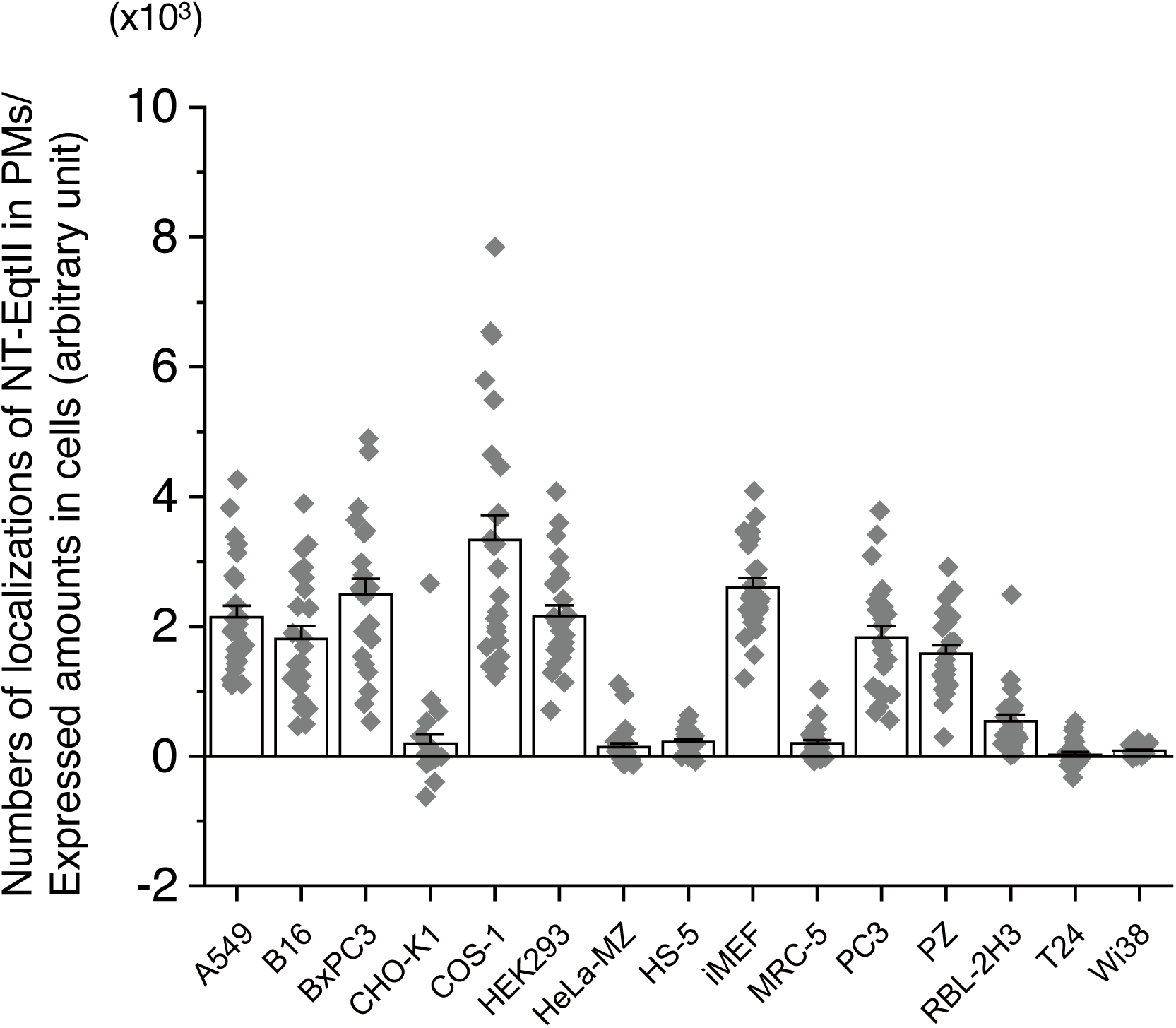
Sphingomyelin is present in the cytosolic leaflets of various cell PMs. The numbers of localizations of NT-EqtII-HaloTag7 labeled with SF650B bound to the cytosolic leaflets of cell PMs, were determined by TIRFM at 4 ms/frame for 2000 frames and subsequently normalized by the expressed amounts (fluorescence intensities in the cytoplasm minus those in the background) measured by oblique illumination. The bars indicate standard errors.

### SM formed small domains that were colocalized with cholesterol and a saturated fatty acid-anchored protein

Since SM is regarded as a major component in forming lipid rafts in the extracellular leaflet of the PM, SM might form rafts also in the cytosolic leaflet of the PM. To investigate whether SM forms such domains, we observed single molecules of NT-EqtII-HaloTag7-SF650B on living cell PMs at 4 ms resolution for 2000 frames by photoactivated localization microscopy (PALM). As shown (Fig. 4), NT-EqtII-HaloTag7-SF650B exhibited small domains of about 130 nm in diameter in the cytosolic leaflet of the PM of a variety of cell lines (Fig. 5A, B).

**Fig. 4.**
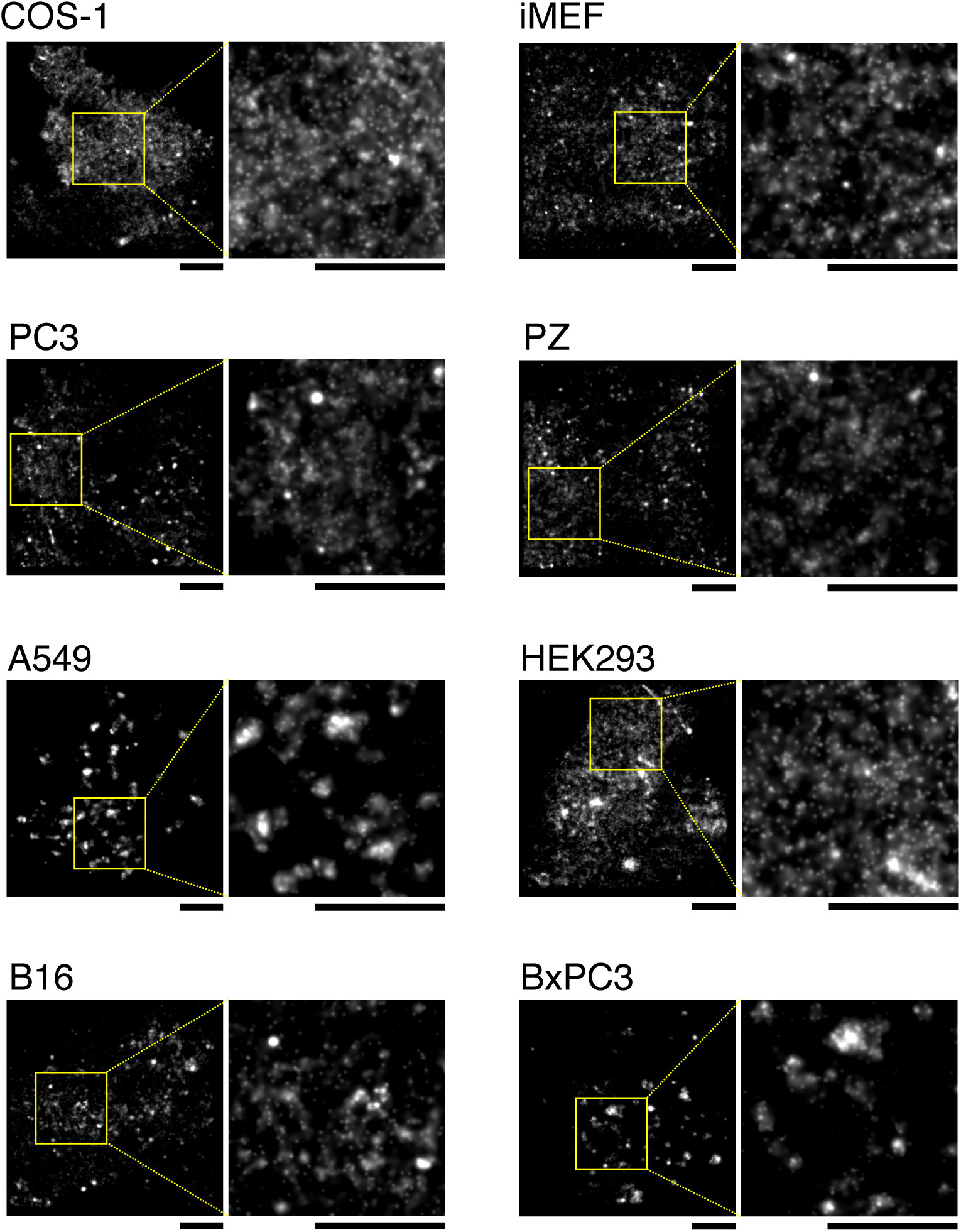
Sphingomyelin forms apparent small domains in the cytosolic leaflets of living cell PMs. The wide field (left) and enlarged (right) dSTORM images of NT-EqtⅡ-HaloTag7 labeled with SF650B in the cytosolic leaflets of PMs of a variety of cells (COS-1, iMEF, PC3, PZ, A549, HEK293, B16, BxPC3). The data acquisitions of dSTORM images were performed at 37°C and at 4 ms/frame for 2000 frames: scale bars, 3 μm.

**Fig. 5.**
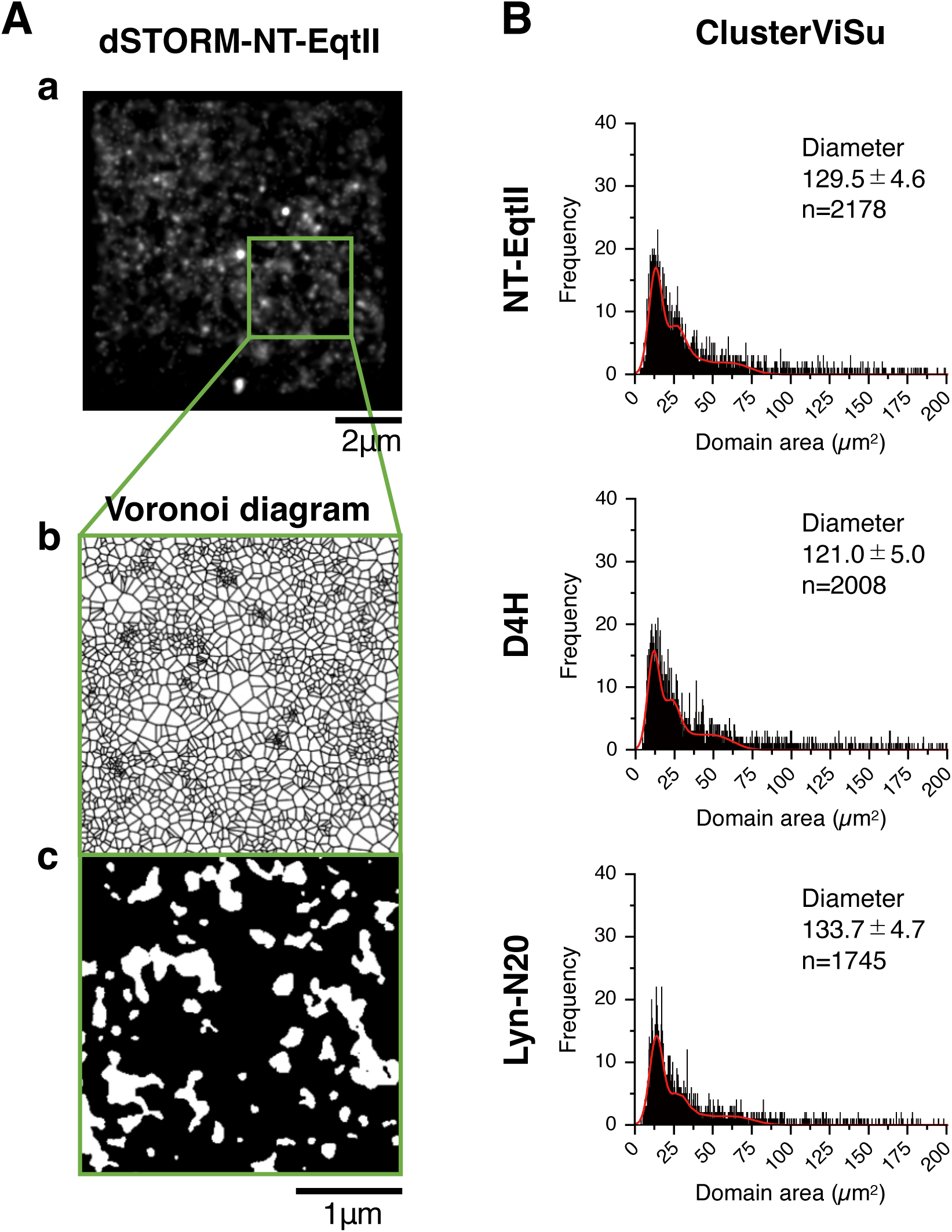
Distributions of domain areas of NT-EqtII, D4H and Lyn-N20 in the cytosolic leaflet of living cell PMs, which were estimated by Voronoi diagram-based analysis algorithms. (A) Typical dSTORM image of (a) NT-EqtII-HaloTag7-SF650B expressed in the living COS-1 cell PM (data acquisition at 4 ms/frame for 2000 frames) and (b) an example of Voronoï polygons by ClusterViSu (Andronov et al., 2016) in the square in (a). (c) The detected domains are shown by white areas. (B) ClusterViSu was used for the estimation of the domain areas. The threshold was automatically determined by Monte Carlo simulation. Since multiple domains at distances less than the spatial resolution are indistinguishable, histograms of the area distribution were fitted with five Gaussian functions to estimate the diameter of the individual domains (Kasai et al., 2011), assuming that the domain to be a circle. The sum of five gaussian curves are shown in red. The average diameters ± standard errors of detected domains are shown in the graphs.

Multi-blinking of dyes for super-resolution microscopy may bring overcounting artifacts (Rossboth et al., 2018), which may lead to a false perception of domain formation. The presence of molecular clusters should be verified by two-color single-molecule localization microscopy (Arnold et al., 2020). Therefore, to verify whether the lipid molecules form domains in the cytosolic leaflet, we next performed simultaneous dual-color observation of stochastic optical reconstruction microscopy (dSTORM) of NT-EqtII and PALM of D4H (a cholesterol probe)/Lyn-N20 (a representative raftophilic molecule) in live COS-1 cells. dSTORM images of NT-EqtII-HaloTag7-SF650B appeared to be partially colocalized with PALM images of mEos4b-D4H or Lyn-N20-mEos4b (Fig. 6A). The average sizes of domains of D4H and Lyn-N20 were comparable to that of NT-EqrtII (approximately 130 nm in diameter) (Fig. 5A, B).

**Fig. 6.**
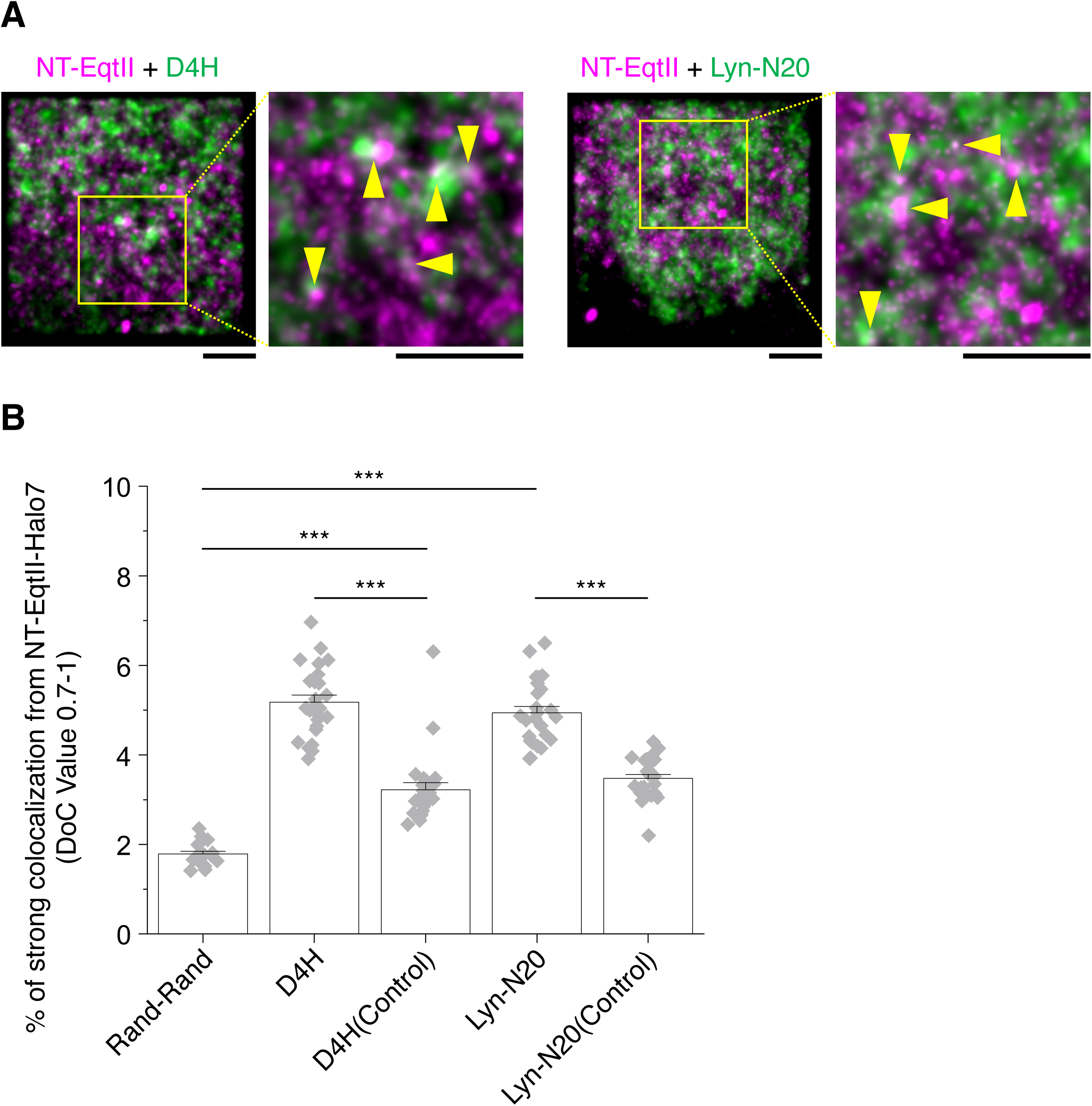
Dual-color super-resolution observation and colocalization analysis between NT-EqtII and cholesterol or Lyn-N20 in living COS-1 cell PMs. (A) Dual-color super-resolution images of NT-EqtII-HaloTag7 labeled with SF650B (dSTORM, magenta) and D4H or Lyn-N20 labeled with mEos4b (PALM, green) of COS-1 cells. The data acquisitions of PALM/dSTORM images were performed at 37°C and 4 ms/frame for 2000 frames: scale bars, 2 μm. Colocalization between domains can be observed as indicated by yellow arrowheads. (B) Degree of colocalization (DoC) scores (0.7-1.0) of NT-EqtII-HaloTag7 and mEos4b-D4H or Lyn-N20-mEos4b. Rand-Rand indicates the DoC value between computer-generated completely random distributions. The control DoC values were obtained by calculating the localization coordinates of NT-EqtII-HaloTag7-SF650B and pseudo-localization coordinates of mEos4b-D4H or Lyn-N20-mEos4b which were generated by shifting the localizations in random directions by random distances. The bars indicate standard errors. *** indicates significant differences (p<0.001) using Welch’s T-test.

We then analyzed the degree of colocalization (DoC) (Malkusch et al., 2012; Pageon et al., 2016) between these domains. The quantitative DoC analysis showed that the localization coordinates between NT-EqtII-HaloTag7-SF650B and mEos4b-D4H or between NT-EqtII-HaloTag7-SF650B and Lyn-N20-mEos4b were distributed in higher DoC scores (0.7-1.0) more frequently than control (Fig. 6B). The control DoC values were obtained by calculating the localization coordinates of NT-EqtII-HaloTag7-SF650B and pseudo-localization coordinates of mEos4b-D4H or Lyn-N20-mEos4b, which were generated by shifting the localizations in random directions by random distances. These results indicated that SM formed small clusters, in which raftophilic molecules such as cholesterol and Lyn-N20 were enriched, in the cytosolic leaflet of the PM in living COS-1 cells.

### Single-molecule imaging at 4 ms resolution showed that NT-EqtII underwent apparent fast simple Brownian diffusion in the cytosolic leaflet of the PM and was not confined within small domains

We next examined whether NT-EqtII, D4H, and Lyn-N20 are temporarily confined in small domains in the cytosolic leaflet of the PM in COS-1 cells. These probes were tagged with tandem dimer (td)-StayGold (Hirano et al., 2022) and analyzed by single fluorescent-molecule imaging. The time resolution was also set at the same (4 ms) as the PALM observation.

Plots of the mean square displacement (MSD)-time revealed that both averages of the diffusion coefficients of the single molecules in the time window of 12 ms and the ensemble-averaged diffusion coefficients on the time scale between 8 and 40 ms were more than 0.8 μm^2^/s at 37°C (Fig. 7; Fig. S2). These diffusion coefficients were comparable to those of a PS probe (2xPH) (Uchida et al., 2011) and a PI(4,5)P_2_ probe (2xPH domain of PLCδ) (Fig. 7; Fig. S2). The statistical analysis showed that almost all these trajectories were categorized as simple Brownian diffusion (Fig. 7A and 7B). Furthermore, no significant entrapment/immobilization longer than 20 ms within <100-nmΦ domains was detected with any of the lipid marker proteins at 37°C (Fig. 8A, B). These results suggested that, while SM formed small domains of about 130 nm which colocalized with domains of cholesterol and Lyn-N20 in the cytosolic leaflet of the PM of COS-1 cells, these lipids were not confined in the small domains for longer than 20 ms but entered into and went out of the domains very rapidly.

**Fig. 7.**
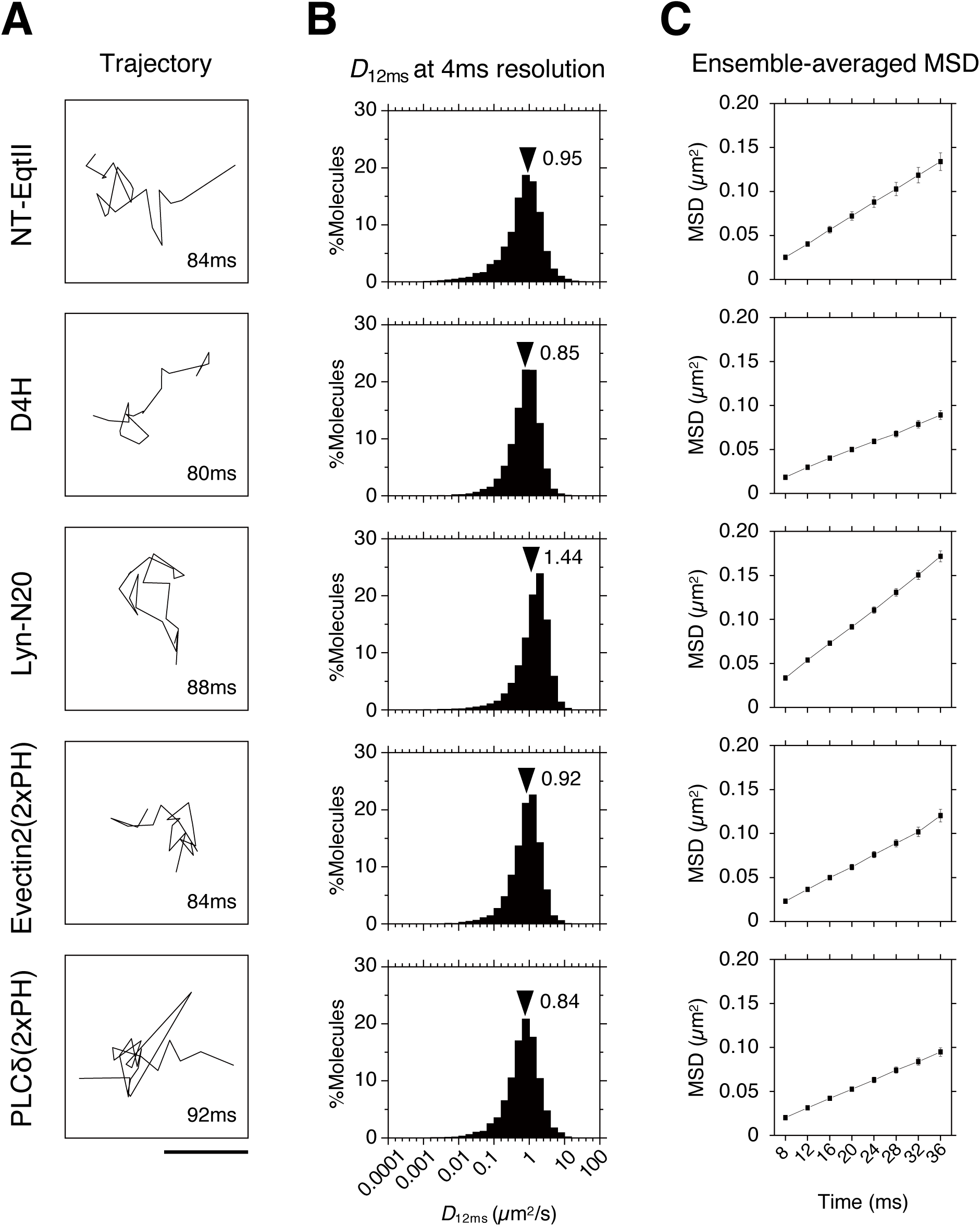
Diffusion of single-molecules of NT-EqtII, D4H, Lyn-N20, evectin 2 (2xPH), and PLC8 (2xPH) at 37 °C, recorded at a time resolution of 4 ms. (A) Typical trajectories of single NT-EqtII, D4H, Lyn-N20, evectin2 (2xPH) and PLC8 (2xPH) fused with tdStayGold in COS-1 cell PMs at 37℃, exhibiting apparent fast simple Brownian diffusion. The lengths of the trajectories are shown at the right-bottom. (B) Histograms showing the distributions of diffusion coefficients of single molecules shown in (A) in the time window of 12 ms. The black arrowheads indicate mean values. (C) Plots of ensemble-averaged mean-square displacements against time (MSD-Δt plots) of the single molecules shown in (A) at 37°C. These plots were practically linear between 8 and 40 ms, suggesting that these molecules underwent effective simple-Brownian diffusion on this time scale. Bars indicate standard errors.

**Fig. 8.**
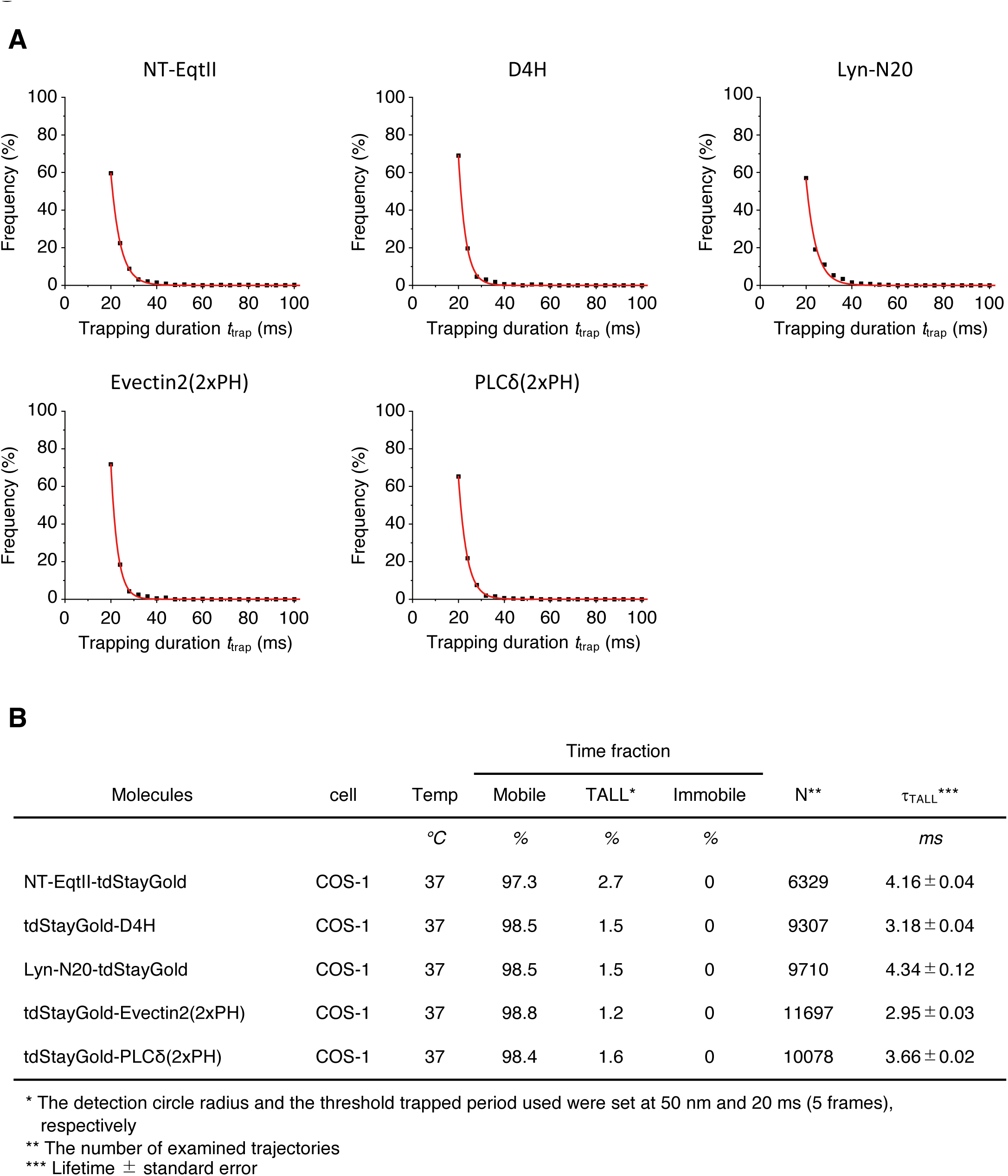
Temporary arrest of LateraL diffusion (TALL) analysis of NT-EqtII, D4H, Lyn-N20, evectin 2 (2xPH), and PLC8 (2xPH) at 37 °C, recorded at 4-ms resolution. (A) Histograms showing the distributions of TALL periods fitted with an exponential decay curve. (B) The time fractions of mobile, TALL, and immobile periods, as well as TALL durations (lifetime, *ι*_TALL_) of NT-EqtII, D4H, Lyn-N20, evectin 2 (2xPH), and PLC8 (2xPH) at 37°C, recorded at 4-ms resolution in the intact COS-1 cell PM. These results show that SM, cholesterol, Lyn-N20, PS, and PI(4,5)P_2_ rarely exhibited TALL in the intact PM. The detection circle radius and the threshold trapped period used were set at 50 nm and 20 ms (5 frames), respectively.

## Discussion

In the present study, we discovered the presence of SM in the cytosolic leaflet of the steady-state PM of various living cell lines for the first time. Furthermore, we performed single-molecule imaging of lipid-binding proteins for SM and cholesterol, and super-resolution single-molecule localization microscopy in the cytosolic leaflet of PMs for the first time.

Bulk SM is synthesized on the lumenal leaflet of the Golgi by SMS1 and transported to the extracellular leaflet of the PM, whereas local SM is synthesized on the extracellular leaflet of the PM by SMS2 (Kumagai and Hanada, 2019; Tafesse et al., 2006). Therefore, the presence of SM in the cytosolic leaflet of biomembranes cannot be explained by the SM synthetic pathway and predicts the molecular machinery that mediates the transbilayer movement of SM from the extracellular (or lumenal) leaflet to the cytosolic leaflet. Indeed, recent studies have identified the molecules involved in the process. PMP2, a causative protein of Charcot-Marie-Tooth disease, induces the tubulation of the PM, which facilitates the flip of SM to the cytosolic leaflet (Abe et al., 2021). TMEM16F, a PM-localized calcium-activated phospholipid scramblase, is indirectly involved in the exposure of the SM to the cytosolic leaflet of lysosomes upon lysosomal membrane damage (Niekamp et al., 2022). In the present study, we found a heterogeneous expression of SM in the cytosolic leaflet in a variety of cell lines. The difference of the expression may give us a clue to identify the molecular machinery that determines the steady-state SM localization in the cytosolic leaflet of the PM.

Simultaneous observation of PALM and dSTORM showed that SM (NT-EqtII) formed small domains that significantly colocalized with small domains of cholesterol (D4H) or Lyn-N20 in the cytosolic leaflet of the PM. On the other hand, single molecules of SM (NT-EqtII) and other lipid-binding proteins showed apparent simple Brownian diffusion in the cytosolic leaflet of the PM when they were observed at 4 ms resolution (Fig. 7), and their diffusion coefficients in the time window of 12 ms and 24 ms were larger than 0.8 μm^2^/s (Fig. 7; Fig. S2). Meanwhile, previous fluorescence correlation spectroscopy observation showed that the diffusion coefficient of the SM probe conjugated with an organic fluorescent dye, ATTO594 via a hydrophilic nonaethyleneglycol (neg) linker in the L_d_ phase of artificial giant unilamellar vesicles was larger than that in the L_o_ phase by a factor of 6-7 (Kinoshita et al., 2017). Therefore, if SM probes temporarily reside in L_o_-like domains in the cytosolic leaflet of the PM, a reduction of the diffusion coefficients and the temporal confinement should be detected. However, SM (NT-EqtII) and other lipid-binding proteins hardly underwent temporal confinement in small domains of 100 nm in diameter (Fig. 8). These results suggested that SM, cholesterol, and Lyn-N20 rapidly diffused in the cytosolic leaflet of the PM, and entered into and went out of the SM-enriched small domains very frequently. The apparent formation of small domains detected by PALM and dSTORM observation may be due to the presence of membrane regions where those lipids are slightly more likely to stay.

Previous studies by ultrafast (25∼100 μs resolution) single-molecule observation (Fujiwara et al., 2002; Fujiwara et al., 2023a; Fujiwara et al., 2023b; Murase et al., 2004; Umemura et al., 2008) have revealed that phospholipid probes exhibited short term (1∼10 ms) confined diffusion in small domains of 40∼230 nm in diameter in the PM, occasionally hopped to an adjacent domain where the phospholipids were again confined. This diffusional behavior is called “hop diffusion”, which can be observed only by ultrafast single-molecule imaging (Fujiwara et al., 2023a; Suzuki et al., 2005) and not by imaging at 4 ms-resolution used in this study. Meanwhile, phospholipids undergo only simple Brownian diffusion in the blebbed PM lacking cortical actin filaments even by ultrafast single-molecule observation (Fujiwara et al., 2002; Fujiwara et al., 2023a). Furthermore, three-dimensional reconstruction of the membrane skeleton by electron tomography (Morone et al., 2006) revealed that the distribution of mesh size of actin-based membrane skeleton located on the PM cytoplasmic surface agreed well with the compartment size distribution determined from the value of phospholipid diffusion, using high-speed single-molecule imaging. Based on these results, it has been proposed that hop diffusion of phospholipids is induced by transmembrane proteins bound to actin-based membrane skeletons, which are called pickets (Kusumi et al., 2005; Suzuki and Kusumi, 2023). The compartment size of phospholipids (DOPE) in COS cells that are used in this study, is reported to be 58 nm on average (Hiramoto-Yamaki et al., 2014). Since the sizes of domains of raftophilic lipids such as SM, cholesterol, and Lyn-N20 observed by PALM and dSTORM were estimated to be approximately 130 nm (Fig. 5A, B) by ClusterViSu method (Andronov et al., 2016), the small domains of these raftophilic lipids would contain at least four compartments formed by the “picket” anchored to the actin-based membrane skeletons. Since amino acid side chains of transmembrane pickets protrude from the α-helix, the rigid tetracyclic ring structure of cholesterol tends to be excluded from the first layer surrounding the transmembrane domain of pockets due to the steric non-conformability between them. This agrees well with the fact that most transmembrane proteins are excluded from the raft-like L_o_-phase domain (Levental et al., 2011). Therefore, the boundary of compartments in which transmembrane picket proteins are aligned would work as the raft breaker, and raft domains may not grow across the compartment boundaries, and the size of rafts would be smaller than the compartment size. This is in line with the fact that large phase separation into L_o_-like and L_d_-like domains is observed in blebbed PMs at low (about 10°C) temperatures, while such large phase separation have never been visualized in intact cell PMs. Nevertheless, the domains of raftophilic lipids of approximately 130 nm in diameter appeared to be formed across at least four compartments made by the transmembrane pickets anchored to the actin-based membrane skeletons (Fig. 9). These results suggest that apparent domains visualized by PALM and dSTORM may consist of assemblies of much smaller domains located within several different compartments (ý=58 nm) made by transmembrane pickets (Fig. 9). Since the spatial resolution of single-molecule localization microscopy is in the range of 20-30 nm (Descloux et al., 2019), it is impossible to distinguish and observe two small domains that are apart from each other at distances less than that. In the future, observation at higher spatial resolution will reveal more precise distribution and colocalization of these lipid domains.

**Fig. 9.**
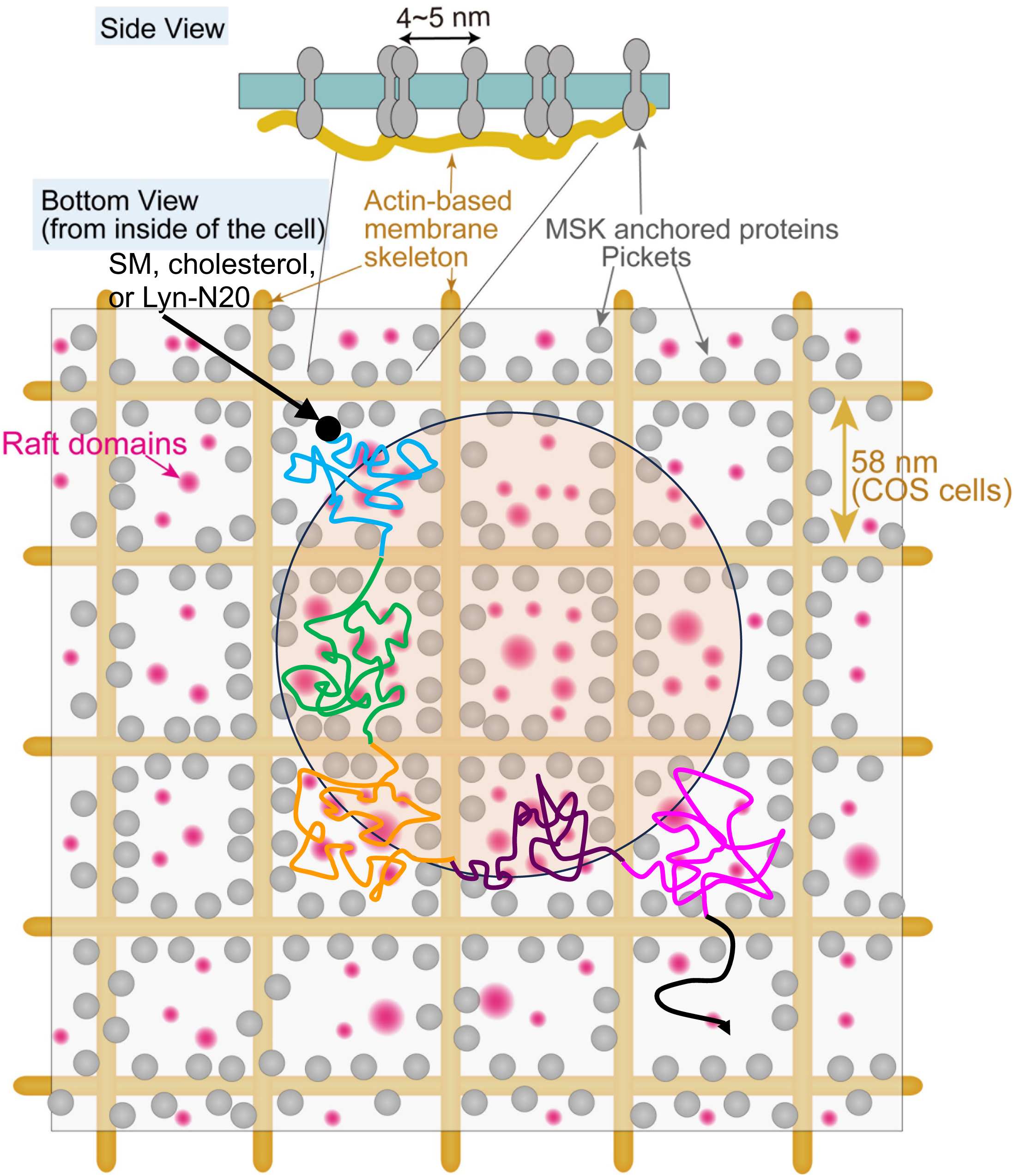
A schematic model for apparent lipid domain formation of about 130 nm in diameter, which were observed by single-molecule localization microscopy. As shown in the bottom view, the cell PM is compartmentalized by raft domains (magenta area) and the transmembrane proteins called “pickets” (gray circle) anchored to the actin-based membrane skeleton. Sphingomyelin (SM), cholesterol, and Lyn-N20 in the cytosolic leaflet of cell PMs are confined within the compartments made by transmembrane pickets due to the steric hindrance and the hydrodynamic friction effects (Fujiwara et al., 2002; Fujiwara et al., 2023a; Fujiwara et al., 2023b) and hop across the compartment boundary to adjacent membrane regions as indicated by the different colors in the trajectory. Since pickets exclude cholesterol, raft domains do not grow across the compartment boundary and the size is smaller than that of the compartment in the cytosolic leaflets of cell PMs. The distance between pickets is estimated to be 4-5 nm according to the Monte Carlo simulation, and the size of rafts that hop across the compartment boundary may be less than 4-5 nm. The apparent domains (beige circle) of approximately 130 nm in diameter that were visualized by single-molecule localization microscopy may consist of assemblies of smaller raft domains located within several different compartments made by pickets.

As mentioned above, the raft-like L_o_ phase would not expand across the compartments made by transmembrane pickets which exclude cholesterol and raftophilic lipids. Montecarlo simulations suggested that the average distance between the transmembrane pickets on the compartment boundary is 4∼5 nm (Ritchie et al., 2005). If SM always resides in moving rafts in the cytosolic leaflets, the rafts hop across the compartment boundaries in which pickets are aligned and need to go through a very narrow corridor. If SM resides in rafts of size larger than the average distance between the pickets, both the hop frequency of the rafts and the diffusion coefficient of SM should be lower than those of non-raft molecules. However, the diffusion coefficients of SM binding protein, NT-EqtII, cholesterol binding protein, D4H and Lyn-N20 were comparable to or even larger than those of PS-binding protein (2xPH of evectin2) and PI(4,5)P_2_ binding protein (2xPH of PLCδ) which possess one unsaturated fatty acid (Fig. 7). These results suggest that the size of the rafts that continue to hold the same SM molecules in the cytosolic leaflets would be very small, presumably less than 4∼5 nm (Fig. 9). Alternatively, if SM frequently enters into and goes out of the moving rafts in the cytosolic leaflets, the size of rafts can be larger than 4-5 nm, but much smaller than the average compartment size of 58 nm. Since the average residency periods of SM, cholesterol, and Lyn-N20 in the compartment of 58 nm were estimated to be 0.85 ms according to (Hiramoto-Yamaki et al., 2014), the residency times of these raftophilic molecules in each raft would be shorter than 0.85 ms (Fig. 9).

We observed the small raftophilic domains in which SM, cholesterol, and Lyn-N20 are temporally colocalized in the cytosolic leaflet of the PM. Lyn-N20 is myristated at Gly2 and palmitoylated at Cys3 (Resh, 1999). Given the affinity of lipidated proteins, in particular, palmitoylated proteins to raft domains (Levental et al., 2011; Lorent et al., 2017), other lipidated proteins that localize at the PM may also have the affinity to the raftophilic domains. The function of the raftophilic domains is currently unclear. However, we envision that the raftophilic domains may serve as the signaling platform, through clustering of signaling PM proteins with lipid modifications in the cytosolic leaflet. Molecular dynamics simulation suggests that very-long-chain (C24) acyl chain in SM interacts with acyl chains of lipids in the opposing leaflet (Rog et al., 2016). Thus, the SM-enriched raftophilic domains in the cytosolic leaflet may be coupled with specific lipid domains in the extracellular leaflet.

In the present study, we developed a novel non-toxic SM probe that allows us to unequivocally demonstrate the presence of SM in the cytosolic leaflet of a variety of living cell PMs. This study also explicitly showed that SM forms apparent small clusters with other raftophilic lipids such as cholesterol and Lyn-N20 in the cytosolic leaflet, where these lipids frequently enter into and go out. This finding would provide a new framework for future studies on signal transduction in the cytosolic leaflet of the PM.

## Materials and Methods

### Plasmids

The plasmids encoding humanized EqtII (Makino et al., 2015) were kindly gifted from Dr. Toshihide Kobayashi. EqtII mutants were generated by site-directed mutagenesis and then introduced into pET45b (+) (Novagen) for bacterial protein expression and pRetroX-TetOne-puro (clontech) for stable gene expression in COS-1 cells. The plasmids encoding bSMase harboring Ras farnesylation sequence [KLNPPDESGPGCMSCKCVLS] (Sakamoto et al., 2019) were kindly gifted from Dr. Yusuf A. Hannun. The cDNA encoding bSMase and Ras farnesylation sequence was subcloned into pEGFP-C3 vector. To generate a catalytically inactive bSMase mutant, D322A/H323A mutation was introduced by site-directed mutagenesis.

To generate an expression vector for NT-EqtII-HaloTag7, the cDNA encoding NT-EqtII was cloned into pEGFP-N1 vector of which GFP was replaced by HaloTag7, the linker sequence 5’-TGGGRASGGGSGG-3’ was inserted between the NT-EqtII and HaloTag7 genes. The cDNA encoding NT-EqtII was also cloned into pPBpuro (puromycin-resistant piggybac-vector). D4H was cloned into pET28/His6-mCherry-D4 (Clone ID: RDB_14300) in which D434S mutation was induced by site-directed mutagenesis (Maekawa and Fairn, 2015). To generate co-expression vector for NT-EqtII-HaloTag7 and mEos4b-D4H, cDNA encoding NT-EqtII-HaloTag7 and mEos4b-D4H were cloned into pEGFP-N1 vector of which GFP was deleted. mEos4b-D4H and EqtII-HaloTag7 were linked by a tandem fusion of two self-cleaving peptides (P2A and T2A), which was previously shown to have higher cleaving activity than the P2A or T2A alone (Liu et al., 2017). To generate expression vector for Lyn-N20, the cDNA encoding Lyn-N20 (Suzuki et al., 2007a; Suzuki et al., 2007b) was cloned into pEGFP-N1 vector, and GFP was replaced by mEos4b. The linker sequence 5’-SGGGGSGGGGSGGGG-3’ was inserted between Lyn-N20 and mEos4b.

To generate expression vector for experiments of single-molecule observation, cDNA encoding 2xPH domain of PLCδ (a kind gift from Dr. Toshiki Itoh, Kobe University) (Furutani et al., 2006), 2xPH domain of Evectin2, Lyn-N20 or D4H was cloned into pEGFP-N1 vector, and GFP was replaced by tandem dimer (td)-StayGold (Clone ID: RDB_19609) (Hirano et al., 2022). The linker sequence 5’-GGGGSGGGGSGGGGS-3’ was inserted into the membrane molecules and tdStayGold.

### Reagents

Bacterial SMase (from Bacillus cereus, S7651) was purchased from SIGMA. Doxycycline (Dox, 631311) was purchased from Clontech. Lipids (egg SM and egg PC) were purchased from Avanti polar lipids as described (Makino et al., 2017; Makino et al., 2015). D609 (CS-0078) was purchased from CEM.

### Cell culture

Immortalized mouse embryo fibroblast cells (iMEFs) and SMS1/2-DKO iMEFs that were kind gifts from Dr. Toshiro Okazaki (Asano et al., 2012), COS-1 African green monkey kidney cells (American Type Culture Collection), and A549 human lung cancer cells were cultured in DMEM supplemented with 10% heat-inactivated fetal bovine serum (FBS) and 1% penicillin/streptomycin/glutamine (PSG) in a 5% CO_2_ incubator. COS-1 cells that could stably express Dox-inducible EGFP, or Eqt-EGFP were established using retrovirus transfection: HEK293T cells were transfected with pRetroX-TetOne-puro contsructs together with pCG-VSV-G and pCG-gag-pol and the medium containing the retroviral particles was collected. COS-1 cells were then incubated with the medium and selected with 2.5 µg/mL puromycin for a week. Chinese hamster ovary (CHO-K1) cells, PC3 human prostate cancer cells (American Type Culture Collection) were maintained in HAM’s F12 medium supplemented with 10% heat-inactivated FBS and 1% penicillin/streptomycin/glutamine (PSG. PZ human prostate cells (American Type Culture Collection) were grown in keratinocyte serum-free medium supplemented with 0.05 mg/ml bovine pituitary extract and 5 ng/ml human recombinant epidermal growth factor (Gibco). T24 human bladder cancer cells were cultured in HAM’s F12 medium supplemented with 10% heat-inactivated FBS and 1% penicillin/streptomycin/glutamine (PSG). Rat basophilic leukemia 2H3 (RBL-2H3) cells were maintained in Eagle’s minimal essential medium supplemented with 1 mM sodium pyruvate, 0.1 mM MEM non-essential amino acids, 15% FBS, and 1% penicillin/streptomycin/glutamine (PSG). MRC-5 human fetal lung fibroblast cells (American Type Culture Collection), HeLa MZ human uterus cervix cancer cells were cultured in DMEM supplemented with 10% FBS and 1% penicillin/streptomycin/glutamine (PSG). WI-38 human lung fibroblast cells (American Type Culture Collection) were cultured in MEM supplemented with 10% heat-inactivated FBS and 1% penicillin/streptomycin/glutamine (PSG). B16 mouse melanoma cells and HS-5 human marrow stromal cells (American Type Culture Collection) were cultured in DMEM supplemented with 10% heat-inactivated FBS and 1% penicillin/streptomycin/glutamine (PSG). BxPC-3 human pancreas cancer cells were cultured in RPMI-1640 medium supplemented with 10% heat-inactivated FBS and 1% penicillin/streptomycin/glutamine (PSG).

### Plasmid transfection

Cells were transiently transfected with plasmids using Lipofectamine 2000 (Invitrogen) or Lipofectamine LTX (Invitrogen) or Lipofectamine 3000 (Invitrogen) or 4-D nucleofector (LONZA) according to the manufacturer’s instructions.

### Expression and purification of recombinant proteins

WT or mutant EqtII cDNA was subcloned into pET45b (+) (Novagen), and the N-terminal 6xHis-tagged and C-terminal EGFP-tagged proteins were expressed in Arctic Express (DE3) RP cells (Agilent Technologies). Bacterial cultures in LB medium were induced with 0.1-0.2 mM IPTG at exponential growth phase (OD_600_ = 0.5-0.6) and grown for 24 h at 11-12°C. The proteins were purified using a HisTrap HP column (GE Healthcare) according to the manufacturer’s instructions, and then dialyzed against PBS using Vivaspin 20 (GE Healthcare).

### ELISA measurement of His-EqtII-EGFP binding to lipids

ELISA was performed as described previously (Makino et al., 2015). In brief, 50 µL of lipid (10 mM) in ethanol was added to each well of a microtiter plate (Immulon 1; Dynatech Laboratories, Alexandria, VA, USA). After solvent evaporation at room temperature, 100 µL of 30 µg/mL bovine serum albumin (BSA) in Tris-buffered saline (TBS; 10 mM Tris-HCl, pH7.4, 150 mM NaCl) was added to each well. After washing, the wells were incubated with 50 µL each of His-EqtII-EGFP protein solution at indicated concentrations in TBS containing 10 µg/mL BSA for 1 h at room temperature. The bound proteins were detected by incubating with peroxidase-conjugated streptavidin. The intensity of the color developed with o-phenylenediamine as a substrate was measured with a Molecular Devices Spectra Max M2 (absorption at 490 nm).

### Measurement of hemolysis

Hemolytic activity was measured in mouse (C57bl/6j) erythrocytes, as described (Makino et al., 2017; Yamaji-Hasegawa et al., 2003). Briefly, erythrocytes were collected by centrifugation, diluted into 1 × 10^7^ cells/mL, and then the 100 µL cell suspension was mixed with each His-EqtII-EGFP protein at the indicated concentrations for 30 min. Ab600 was measured by a plate reader.

### bSMase treatment

Live COS-1 cells were treated with 1 Unit/mL bSMase in serum-free DMEM for 60 min at 37°C as described previously (Yachi et al., 2012).

### LDH release assay and MTT assay

Viability of COS-1 cells (1.2 × 10^4^ cells/well) was measured with MTT (3-(4,5-dimethylthial-2-yl)-2,5-diphenyltetrazalium bromide) by using Cell Counting Kit-8 (Doujin) according to the manufacturer’s protocol (Makino et al., 2017). The release of LDH from COS-1 cells (3.2 × 10^3^ cells/well) was measured by using Cytotoxicity LDH Assay Kit-WST (Doujin) according to the manufacturer’s protocol (Makino et al., 2017).

### Immunocytochemistry

Cells were washed with PBS briefly, fixed with 4% Paraformaldehyde in PBS at room temperature for 15 min, and quenched with 50 mM NH_4_Cl in PBS at room temperature for 10 min. Cells were then stained with 1 µg/mL His-EqtII-EGFP and DAPI (D9542, SIGMA) at room temperature for 1 h, washed with PBS, and then mounted in PermaFluor (Thermo).

### Confocal microscopy

Confocal microscopy was performed using a TCS SP8 (Leica) with a 63 x 1.2 Plan-Apochromat oil immersion lens. Excitation was performed with a 65-mW argon laser emitting at 405 nm (for DAPI) or 488 nm (for GFP). Emissions were collected at 420– 480 nm (for DAPI) or 500–600 nm (for GFP) using a Spectral detector.

### Fluorescent image analysis

Quantitation of images was performed with ImageJ software (NIH).

### Propidium iodide (PI) staining and detection by flow cytometry

Expression of EGFP or EqtII-EGFP was induced in COS-1 cells after Dox-treatment for the indicated times. Cells were then collected, stained with PI for 15 min at RT, and kept on ice. Fluorescence of EGFP and PI was measured by FACS Calibur (BD Biosciences).

### Quantification of NT-EqtII in the cytosolic leaflet of the PM

The quantities of NT-EqtII-HaloTag7, labeled with SF650B and expressed in the cells were quantified by measuring the fluorescence intensities of the molecules observed by oblique illumination (Fig. 2B, left). Subsequently, the fluorescence intensities of cells in the images stacked 2000 frames were determined using ImageJ, with the subtraction of fluorescence intensity originating from the cell-free region serving as the background. Furthermore, the quantification of the NT-EqtII-HaloTag7 contents in the cytosolic leaflet of the cell PM was accomplished by measuring numbers of the single fluorescent spots, using home-built, objective-lens-type total internal reflection fluorescence microscopy (TIRFM) based on Nikon Ti2 inverted microscope (100x 1.49 NA oil objective) equipped with a high-speed gate image intensifier (C9016-02 MLG; Hamamatsu Photonics) coupled to a sCMOS camera (ORCA-Fusion; Hamamatsu Photonics) (Fig. 2B, right). Numbers of the single spots were measured using the ThunderSTORM plugin of ImageJ with “Wavelet filtering” (B-Spline order = 4 and B-Spline scale = 4.0) and the “Local maximum method” (Peak intensity threshold = 30∼50 and Connectivity = 8-neighbourhood). To eliminate non-specifically bound fluorescent molecules, numbers of the single spots in the cytosolic leaflet of PMs from cells lacking NT-EqtII-HaloTag7 expression, but treated with SF650B under the same condition, were subtracted. Subsequently, numbers of the single spots of NT-EqtII-HaloTag7 in the cytosolic leaflet of cell PMs, were normalized by the expressed quantities in the cells. This method was applied to the data in Fig.3 as well.

### Dual-color super-resolution microscopic observation of NT-EqtII, D4H and Lyn-N20 in living cells

For live-cell dual-color PALM and dSTORM observations of NT-EqtII, D4H and Lyn-N20, COS-1 cells were sparsely seeded in a glass-base dish (4×10^3^ cells on the glass window of 12 mm in diameter, 0.15 mm-thick glass; Iwaki), and grown in DMEM with 10% FBS for 2 days before each experiment. Before observation, these HaloTag7-tagged molecules were labeled with SaraFluor650B (SF650B) by incubating the cells with 10nM of the SF650B-conjugated Halo-ligand for 12min. Single-molecules of mEos4b-D4H, Lyn-N20-mEos4b and NT-EqtII-HaloTag7-SF650B were observed using TIRFM based on a Nikon Ti2 inverted microscope (100x 1.49 NA oil objective) equipped with two high-speed gate image intensifiers (C9016-02 MLG; Hamamatsu Photonics) coupled to two sCMOS cameras (ORCA-Fusion; Hamamatsu Photonics). Single-molecule observation in cells was performed by illumination with an activation laser (405 nm, approximately 10.7 nW/mm^2^ for PALM observation) and excitation lasers (561 nm, approximately 28.1 mW/mm^2^ for PALM and 647 nm, approximately 55.7 mW/mm^2^ for dSTORM). In the excitation arm, a multiple-band mirror (Di01-R405/488/561/635-25×36, Semrock) was employed. The two-color fluorescence images of mEos4b and SF650B were separated into the two detection arms of the microscope by a dichroic mirror (Chroma: ZT561rdc-xr-UF2 or ZT647rdc-UF2). The detection arms were equipped with band-pass filters of FF01-600/37-25 (Semrock) or ET700/75 (Chroma). The data acquisitions of PALM/dSTORM were simultaneously performed at 37°C and at 4 ms/frame for 2000 frames with an image size of 288×288 pixels. Each images were cropped to 120×120 pixels for analysis. The detection of the fluorescent spots in the images was performed by using the ThunderSTORM plugin of ImageJ with “Wavelet filtering” (B-Spline order = 4 and B-Spline scale = 4.0) and the “Local maximum method” (Peak intensity threshold = 30∼50 and Connectivity = 8-neighbourhood). After spot detection, the post-processing steps of “Remove duplications” (Distance threshold = uncertainty) and “Drift correction” (cross-correlation with 5 bins) were further performed. To eliminate the effect of difference of localization densities, 15,000 localizations per image were randomly acquired for all the observations. The superimposition of images in different colors obtained by two separate cameras was performed as reported previously (Komura et al., 2016; Suzuki et al., 2007a; Suzuki et al., 2007b; Suzuki et al., 2012). The single-molecule localization precision of mEos4b and SF650B was 26.3 nm and 22.0 nm, respectively. The final magnifications were 2×, resulting in pixel sizes of 78 nm (square pixels).

### Estimation of domain sizes by ClusterViSu

To estimate the areas and diameters of domains of NT-EqtII, D4H and Lyn-N20, we performed Voronoi-based image segmentation and cluster analysis, called ClusterViSu (Andronov et al., 2016). This method is more suitable than the SR-Tessler method (Levet et al., 2015) in case that clustered points have weak density and shows a more complete detection of the cluster numbers and retrieval of their size and homogenous shape. To preserve detection accuracy, clusters containing five localizations or less were eliminated according to the previous report (Andronov et al., 2016). Since multiple domains at distances less than the spatial resolution are indistinguishable, histograms of the area distribution were fitted with five Gaussian functions to estimate the diameter of the individual domains (Kasai et al., 2011), assuming that the domain to be a circle. The fitting was performed using OriginPro 2018b (OriginLab), assuming the sum of five gaussian functions,

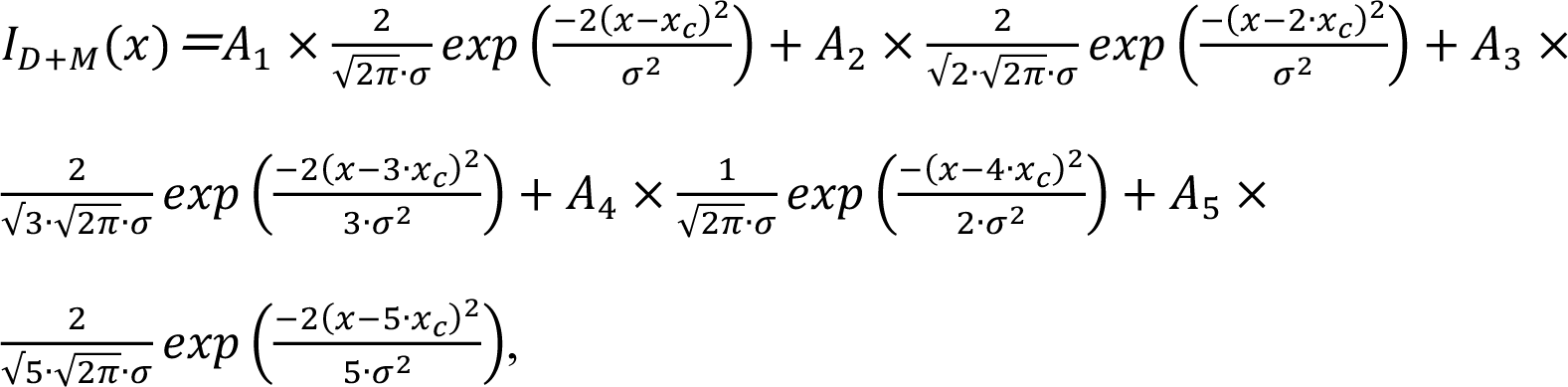

where *x_c_* is the mean value and *0* is the standard deviation. *σ*, *x_c_*, *A_1_, A_2_, A_3_, A_4_* and *A_5_* are the free parameters, and the regression was done by the Levenberg-Marquardt method until χ^2^ reached the minimum.

### Quantitative analysis of the degree of colocalization between NT-EqtII-HaloTag7-**SF650B and mEos4b-D4H or Lyn-N20-mEos4b**

To this end, we performed degree of colocalization (DoC) analysis of PALM and dSTORM data according to (Malkusch et al., 2012) with modification. In brief, to estimate DoC value, for each molecule of protein A, the number of localizations of protein A (NT-EqtII-HaloTag7-SF650B) and protein B (mEos4b-D4H or Lyn-N20-mEos4b) within circles of the increasing radius was calculated, respectively, providing the density gradients of localizations of protein A and protein B around these molecules.

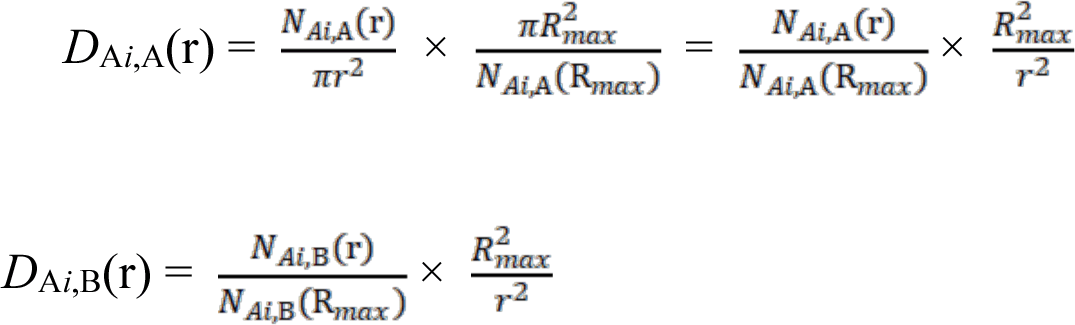

Here, *N*_A*i*,A_(r) is the number of localization of protein A within the distance r around protein A*i*, and *N*_A*i*,B_(r) is the number of localization of protein B within the distance r around A*i*. Then, these density gradients were corrected for the area (πr^2^), normalized by the number of localizations within the largest observed distance R*_max_* and divided by the largest area for protein A 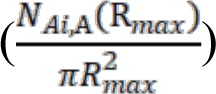 and protein B 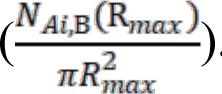. Namely, the density gradients were corrected by the density at the maximum radius respectively for protein A and protein B. R*_max_* and dR (the bin of radius for analysis) were set at 500 nm and 50 nm, respectively, and if both the number of localizations of protein A and that of protein B were less than 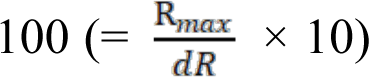 in a circle with R*_max_*, CBC values were not calculated because the density gradients cannot be accurately calculated. A uniform distribution gives an expected value of *D*(r) = 1 for all r.

The two distributions were compared by calculating a rank correlation coefficient (Spearman), in which the colocalization coefficient was weighted by a value proportional to the distance to the nearest neighbor to avoid long-distance effects (Malkusch et al., 2012).

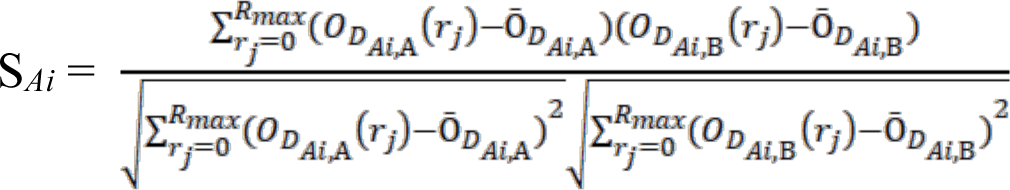

Here, 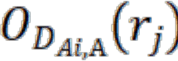 is the rank of 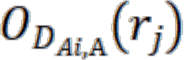 calculated after Spearman, and 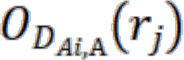 is the arithmetic average of 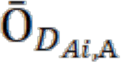. The colocalization value C*_Ai_*, was calculated as C*_Ai_* = S*_ai_* ×, 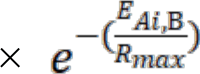, with E*_Ai_*_,B_ as the distance from *Ai* to the nearest neighbor from protein B. C*_ai_* (CBC score) was calculated for every single-molecule localization and ranged from -1 (anti-colocalized or segregated), through 0 (no colocalization), to +1 (totally colocalized). Furthermore, C*_Bi_* was also calculated as well. The summation of C*_Ai_* and C*_Bi_* in the bins between 0.7 and 1 was used as an index for the colocalization. For control analysis, PALM images of mEos4b-D4H or Lyn-N20-mEos4b were replaced by pseudo-localization coordinates which were generated by shifting the localizations in random directions by random distances and overlapped with dSTORM images of NT-EqtII-HaloTag7-SF650B and the DoC scores were estimated.

### Single fluorescent-molecule tracking

For single fluorescent-molecule video imaging, COS-1 cells expressing NT-EqtII-tdStayGold, Lyn-N20-tdStayGold, tdStayGold-D4H, tdStayGold-2xPH (PLC8), or tdStayGold-2xPH (evectin2) were sparsely seeded in a glass-base dish (4×10^3^ cells on the 12-mm diameter glass window, 0.15-mm-thick glass; Iwaki), and grown for 2 days before each experiment.

Single molecules of these lipid-binding proteins or lipid-anchored proteins were observed at 37 °C at 4 ms/frame, using the Nikon Ti2 inverted microscope (100×1.49 NA oil objective) mentioned above as reported previously (Kinoshita et al., 2017; Komura et al., 2016; Morise et al., 2019). The precision of the position determinations for single stationary fluorescent tdStayGold probes was estimated to be 22.2 nm from the standard deviations of the determined coordinates of the probes fixed on coverslips.

Diffusion coefficients for individual spots were obtained as previously described (Kinoshita et al., 2017; Suzuki et al., 2005). In brief, the single-molecule mean square displacement (MSD for the time interval *Δt*, *i.e.*, *ýr*(*ýt*)^2^) is defined as follows. For a single-molecule trajectory consisting of N determined coordinates (*x*-, *y*-positions) in a two-dimensional plane, all of the [*N*-*n*+1] partial trajectories of *n* consecutive positions (*n* ≤ *N*) were extracted. The MSD (*N*, *n*) was then calculated by averaging the square displacements of n steps for all of these [*N*-*n*+1] partial trajectories and, by varying *n*, the plot of MSD (ý*t*_n_ = *n*8*t*) versus n8t (*8t* = the duration of each image frame) was obtained. Namely, the MSD for every time interval was calculated according to the following formula:

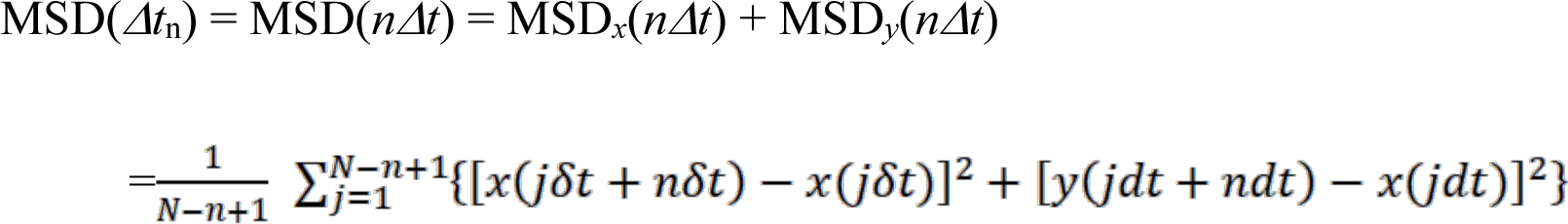

where dt is the frame time, *x*(*j8t +n8t*) and *y*(*j8t +n8t*) describe the particle position following a time interval, *ýt*_n_ = *n8t*, after starting at position (*x*(*jδt*), *y*(*jδt*)), *N* is the total number of frames in the recording sequence, and *n* and *j* are positive integers (*n* determines the time increment). The apparent simple-Brownian diffusion was characterized by a single parameter, the “effective diffusion coefficient”. Namely, under limited conditions of the time resolution and the analysis time window, the MSD-Δt curve can be linearly fit, and only under these circumstances, the diffusion can “effectively” can be described by an “effective diffusion coefficient”, given by the slope of the plot (divided by 4, by definition). The effective diffusion coefficient of a particle in the time window of 12 ms (*D*^eff^_12 ms_) and 24 ms (*D*^eff^_24 ms_) was obtained by linearly fitting its single-molecule MSD-D*t* plot at the 8 ms, 12 ms, 16 ms timepoints and at 8-40 ms timepoints (the slope divided by 4 gives the diffusion coefficient).

### Detection of temporal confinement in nanoscale domains

We attempted to detect temporary confinement/binding events or Temporal Arrest of LateraL diffusion (TALL) events, using the methods developed by (Sahl et al., 2010), with our previous modifications (Kinoshita et al., 2017; Komura et al., 2016). Trajectories longer than 10 frames were used for the analysis (more than 7238 trajectories for all the molecules, with a total number of frames for each molecule greater than 72380). The detection circle radius and the threshold residency time were set at 50 nm or smaller than 10 frames (4 ms), respectively, according to (Sahl et al., 2010). tdStayGold fixed on a glass surface was used as the standard for immobile molecules. The results obtained by this method were comparable to those previously reported by (Simson et al., 1995) and our lab (Kinoshita et al., 2017; Komura et al., 2016; Suzuki et al., 2007a; Suzuki et al., 2007b; Suzuki et al., 2012).

### Statistical analysis

Error bars displayed throughout this study represent s.d. unless otherwise indicated and were calculated from triplicate samples. Data shown are representative of three independent experiments, including microscopy images and western blotting.

## Acknowledgements

We would like to thank Shinobu Kawaguchi for constructing various cDNAs, Yumi Matsuno for assisting with the cell culture, Rinshi Kasai and Koichiro M. Hirosawa for development of the microscope system.

## Footnotes

## Author contributions

T.M., T.N. and Y.U. designed and performed the experiments, analyzed the data, interpreted the results and wrote the paper; K.M. and Y.K. designed the experiments, analyzed the data, interpreted the results, and wrote the paper; T. Kishimoto designed and performed the experiments, analyzed the data, interpreted the results, and wrote the paper. A.M. performed the experiments and analyzed the data; T. Kobayashi and H.A. discussed the results; Y.Y. developed analysis software; T.T. and K.G.N.S. designed the experiments, interpreted the results, and wrote the paper.

## Data availability

All relevant data can be found within the article and its supplementary information.

---

**Fig. S1.**
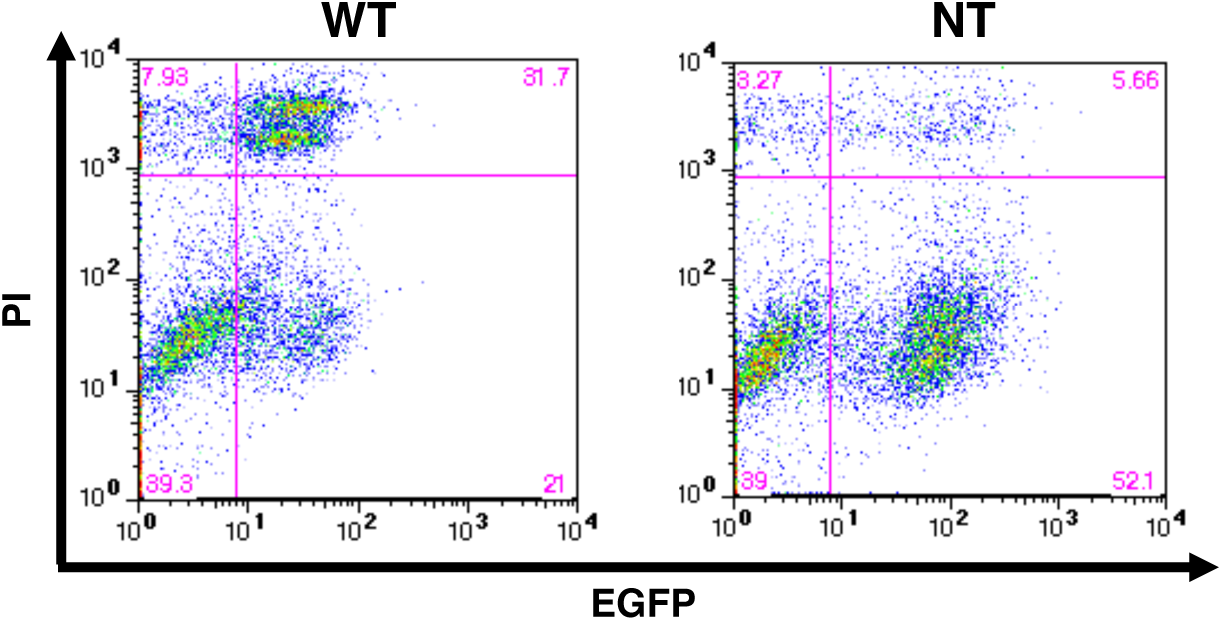
Cytotoxicity of COS-1 cells expressing WT or NT-EqtII-EGFP in the cytosol. COS-1 cells that stably express WT-EqtII-EGFP or NT-EqtII-EGFP in the cytosol in a doxycycline (Dox)-inducible manner were treated with Dox at 1 µg/mL for 45 h. Cells were stained with propidium iodide (PI) and analyzed with flow cytometry. The percentages of cells sorted into each section are shown at the corners.

**Fig. S2.**
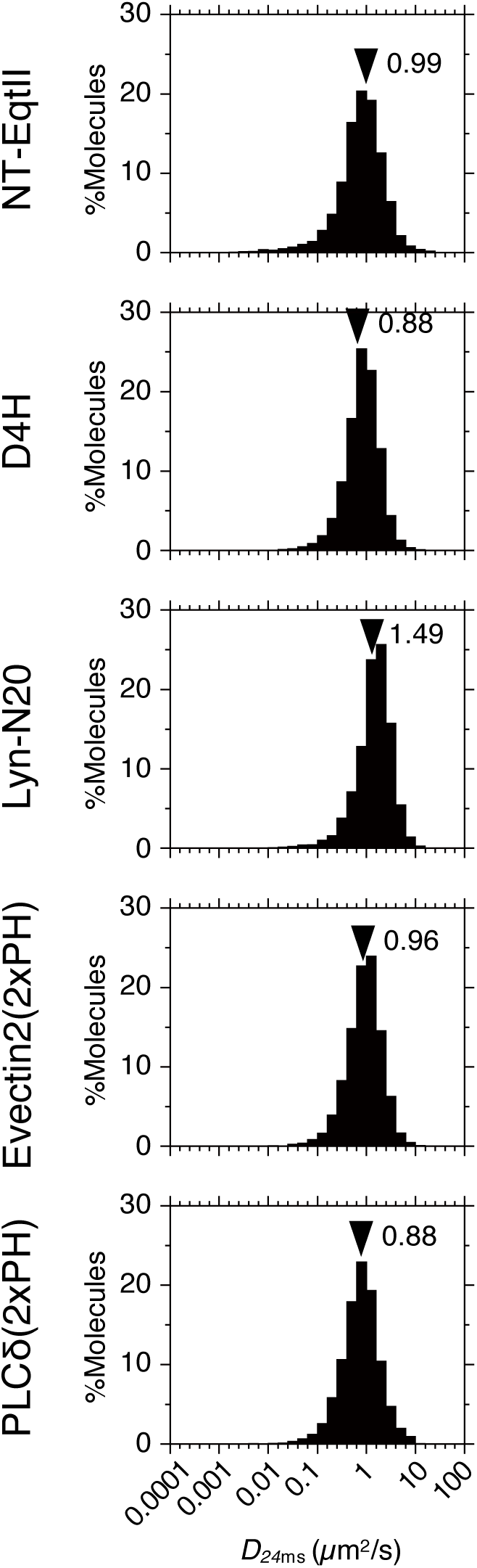
Diffusion coefficients of NT-EqtII, other molecules in the cytosolic leaflet of the living cell plasma membrane. Single molecules of NT-EqtII, D4H, Lyn-N20, Evectin2(2xPH) and PLCδ(2xPH) tagged with (td)-StayGold in the cytosolic leaflet of the COS-1 cell plasma membranes were observed at 4-ms resolution and 37°C. Diffusion coefficients (D_24ms_) were evaluated from the slope of plots of mean square displacements against time (MSD-τιt plots) between 8 and 40 ms. Black arrowheads indicate the mean values of D_24 ms_.

